# Cryo-Correlative Light and X-ray microscopies: Expanding the Intracellular Chemical Map

**DOI:** 10.1101/2025.05.23.655741

**Authors:** D. Karpov, L. Cuau, R. Shishkov, C. Gramaccioni, E. Dallerba, B.J. Schwehr, M. J. Hackett, S. Plush, M. Massi, F. Lerouge, P. Cloetens, S. Bohic

**Author notes:** Correspondence should be addressed to: S.B. or P.C.

## Abstract

We introduce a powerful, integrated workflow that fuses cryo-optical fluorescence microscopy with cryogenic synchrotron radiation X-ray fluorescence nanoimaging to unlock unprecedented nanoscale insights into cellular ultrastructure and composition. Our method delivers sharp 2D and 3D visualizations enabling simultaneous elemental mapping, nanoparticle tracking, and imaging of mitochondrial features via a luminescent cyclometalated iridium complex. We further demonstrated that combining well-chosen molecular probes possessing different heavy elements (e.g. rhenium, iridium, bromine and iodine) allows elemental multiplex “painting” of different organelle to provide X-ray fluorescent elemental contrast of some intracellular structure. By eliminating the need for separate sample preparations, this streamlined approach maximizes limited synchrotron beamtime and dramatically accelerates data acquisition, setting a new benchmark for advanced cryo-nanoscale imaging studies.

## Main text

Biological organisms exhibit complex functionality across various hierarchical levels, from organs and tissues to individual cells and their organelles. The interplay between chemical composition, morphology, and organelle function is critical for understanding biochemical processes in both healthy and diseased cells. Understanding these functionalities requires the characterization of the processes and their underlying properties at different scales. The integrative study of the cellular and molecular process has only just become possible as modern imaging methods recently achieved the required levels of resolution and sensitivity through continuous improvement in light and electron microscopy. The recent advent of light fluorescence nanoscopy has opened up particularly stunning possibilities [1]. Despite that, it still provides little to no chemical information about the subcellular compartments – essential information for understanding the localization and function of cellular machinery.

The cell ionome [2] is a crucial characteristic that influences pathophysiological and toxicological processes [3-6]. This is especially important as humans face increasing exposure to exogenous compounds such as metal nanoparticles and metallic pollutants, alongside changes in trace element homeostasis. These factors define the human exposome and have significant, yet largely unexplored, health implications. Moreover, quantitative mapping of elemental distributions aligns with overarching goals in modern analytical chemistry of understanding the role of nanoscale chemical heterogeneities in complex processes.

Synchrotron-based X-ray nanoprobes provide a powerful approach to the cell ionome studies, down to the organelle^3^ and sub-organelle level, where the local structural variations within organelles can uncover new ways to characterize cellular taxonomy [7]. These methods comprise sequential imaging by cryogenic soft X-ray microscopy [8] to obtain 3D high-resolution ultrastructural information, followed by cryogenic synchrotron radiation X-ray fluorescence nanoimaging (SR-XRF-N) of the same cryo-preserved cell, thereby providing complementary elemental mapping. Importantly, the elemental composition of most organelles is relatively uniform and does not significantly differ from that of the surrounding cytoplasm, and often fail to produce measurable contrast in X-ray fluorescence images. At room temperature, a quasi-correlative approach that combined SR-XRF-N with transmission electron microscopy (TEM) on very thin cellular sections, allowing for high-resolution structural and elemental analysis simultaneously [9]. However, synchrotron-based nanoprobe techniques often fall short of achieving desirable statistical significance due to constraints on sample throughput, and the reliance on synchrotron facilities – available only through scheduled access – which limits their continuous applicability compared to laboratory-based systems.

Here, we present a combination of high-throughput laboratory-based cryo-optical fluorescence light microscopy (cryo-FLM) to pre-screen and prioritize samples, and advanced cryogenic SR-XRF-N developed and integrated at the ID16A nano-imaging beamline of the new Extremely Brilliant Source (EBS) at the European Synchrotron Radiation Facility (ESRF).

We developed a workflow that ensures reliability and improved identification of the intracellular target compartments in vitrified cells, when using molecular probes that possess a heavy element (e.g. transition metal) (Fig. 1a-f). The sample storage and transfer are designed to keep samples below 120K throughout the investigation (Fig.1, Supplementary Fig. 1). Our cryo-FLM, enhanced with computational optical-clearing, delivers detailed subcellular imaging of the labelled organelle for all cells grown on Si_3_N_4_ membrane support (Fig. 1f). Recent improvements in X-ray detection efficiency have been achieved by adding a prototype 16-element SDD Ardesia detector [10]. Furthermore, improved stability and acquisition schemes through the implementation of interferometers has allowed for high-sensitivity elemental mapping which extends the proposed correlative cryo-workflow for 3D elemental nanotomography (Fig. 1k).

**Figure 1:**
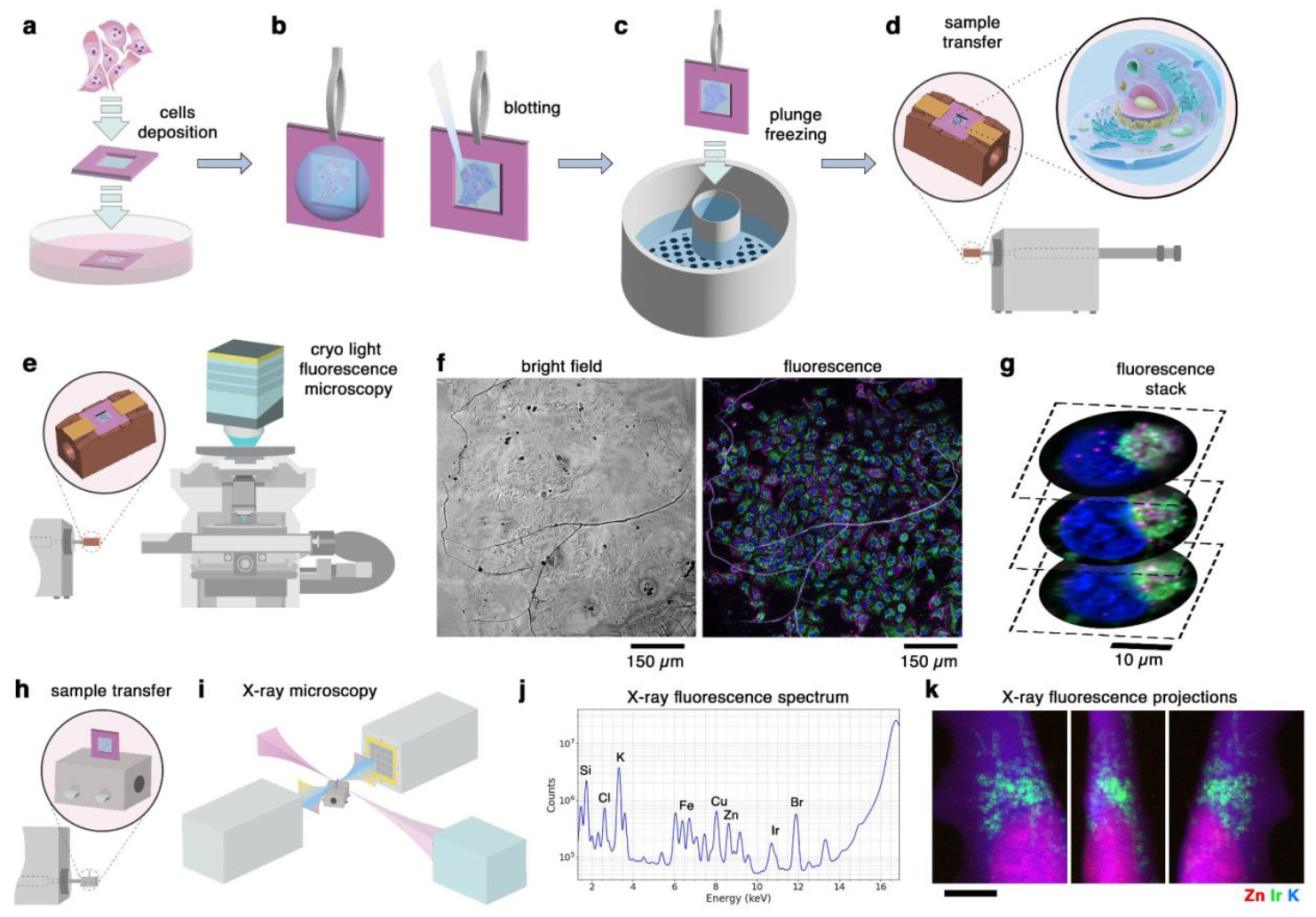
Cryogenic workflow for 2D/3D Cryo-correlative optical fluorescence microscopy and X-ray elemental nanoimaging of entire frozen-hydrated cell. **(a)** Cells are directly grown as a monolayer onto small 500 nm-thick Si3N4 membrane coated with a few atomic layers of graphene. **(b & c)** The cell monolayer is rinsed in a physiological ammonium acetate buffer followed by a double-side manual blotting prior immediate plunge freezing in liquid ethane at 93K resulting into cryogenically immobilized cell fixed in a vitreous state preserving ultra-structural and chemical integrity of the cell whilst vitrified sample kept at LN2 temperature allows a slowing-down of radiation damages under intense X-ray focused beam. **(d)** As such, upon specific unique home-made developments and adaptation to the synchrotron ID16A nanoprobe end-station, dedicated vacuum-based cryogenic sample stage and cryogenically-cooled cryotransfer systems were implemented to keep frozen-hydrated cell monolayer on the Si_3_N_4_ membrane below 120 K. **(e)** For the cryo-FLM sample transfer, a dedicated copper-based cartridge was developed to accommodate Si_3_N_4_ membrane enabling cryo-transfer and observation of the very same vitrified cells. At all steps, sample contaminations, such as hexagonal ice, can occur and should be reduced by minimizing ice contamination using freshly clean and decanted liquid nitrogen and working as far as possible in a low humidity-controlled environments. **(f & g)** Cryo-FLM is performed first and an entire image of the membrane is registered as a mosaic in bright-field imaging mode and followed by a full Z-stack multi-channel fluorescence imaging. The latter is processed through computational clearing and maximum integrated intensity of fluorescence images stack. **(h)** The Si_3_N_4_ membrane is then mounted and clamped vertically within dedicated aluminium-based sample holder in liquid nitrogen and the whole assembly is moved to the docking port of the high-vacuum ID16A endstation using a vacuum cryo-transfer system EM VCT500 holding the samples at 110 K. The bright-field image loaded in a home-made scanning/imaging graphical interface of the end-station allow to define correspondence between the pixel values of the 4 corners of the membrane window and the motor positions using the long-working distance online optical visible microscope of the end-station for positioning of the membrane, in addition a small 0.5 mm orientation membrane made in one corner of the Si frame is of additional help. The calculated homography matrix takes into account the in-plane orientation of the membrane. This further allow to navigate dynamically in the mosaic image either bright-field or fluorescence and select cell of interest or a particular intracellular labelled region of interest for subsequent detailed investigation through cryo-SR-XRF-N raster-scanning **(i)** with pixel-based XRF spectrum acquisition that allow to display the elemental composition of the scanned areas **(j)** resulting in elemental maps. **(k)** The SR-XRF-N enable not only high resolution 2D scanning but also missing-wedge tomographic tilt series acquisition of XRF maps (typically in 3° increments, throughout a tilt range of approximately −70° to +70°) of a targeted cell or cellular region when properly aligned on the rotation axis of the cryo-sample stage assembly. These images are then aligned and computationally merged into a 3D reconstruction of the sample, or tomogram.

We demonstrate the strength of this combinatorial approach by imaging a molecular probe based on a luminescent cyclometataled iridium complex, herein referred to as Ir-Mito. Previous investigations demonstrated that Ir-Mito exhibits specificity for mitochondrial structures in cells [11] (Supplementary Fig. 2). Additionally, we treat the cells with pegylated GdTbF_3_ nanoparticles, that we designed for X-ray theragnostic purposes [12-14]. Our approach allows us to study both the mitochondrial structures highlighted byIr-Mito and the uptake of the pegylated GdTbF_3_ nanoparticles. This elemental multiplexing approach avoids the need for complex sample preparation for separate experiments and ensures that comprehensive results can be obtained within limited experimental time available at synchrotron facilities.

Specific elemental contrast from cell organelles and exposed to GdTbF_3_-pegylated nanoparticles are obtained. The luminescent Ir-Mito complex depicts mitochondria in cryo-FLM that, as expected, correlated very well with the Ir 2D SR-XRF-N projection map (Fig. 2). X-ray phase contrast imaging of the frozen-hydrated cells can also be acquired as part of routine capture prior to XRF scanning with an effective total dose of only few hundred Gray [15]. It allows the cell shape and the contours of the nuclear and nucleoli to be discerned and correlated with the well-preserved elemental distribution detected by XRF, using zinc to delineate the nuclear shape and phosphorus to reveal the nucleoli and nuclear phosphate-rich chromatin regions [16] (Fig. 3; Supplementary Fig. 3). The gadolinium and terbium were representative of the intracellular uptake and location of the GdTbF_3_-PEG nanoparticles after 24h exposure and correlated to the position of fluorescently-labeled lysosomes (Fig. 2, Supplementary Fig. 3 & 4), as intracellular uptake is expected to be most likely through the endocytic route. This will now require further in-depth studies beyond the scope of the present work. The iridium distribution revealed by cryogenic SR-XRF-N clearly delineates mitochondria but unexpectedly internal lamellar structures were also made visible. Lines profile along mitochondria found structures separated in the 60-150 nm range (Fig. 2I & J, Supplementary Fig. 3) that fall in the dimension of cristae structures similar to what is reported using STED-nanoscopy [17]. Such a level of detail, achieved by specifically detecting the iridium X-ray fluorescence from Ir-Mito inside mitochondria, has not been demonstrated until now.

**Figure 2:**
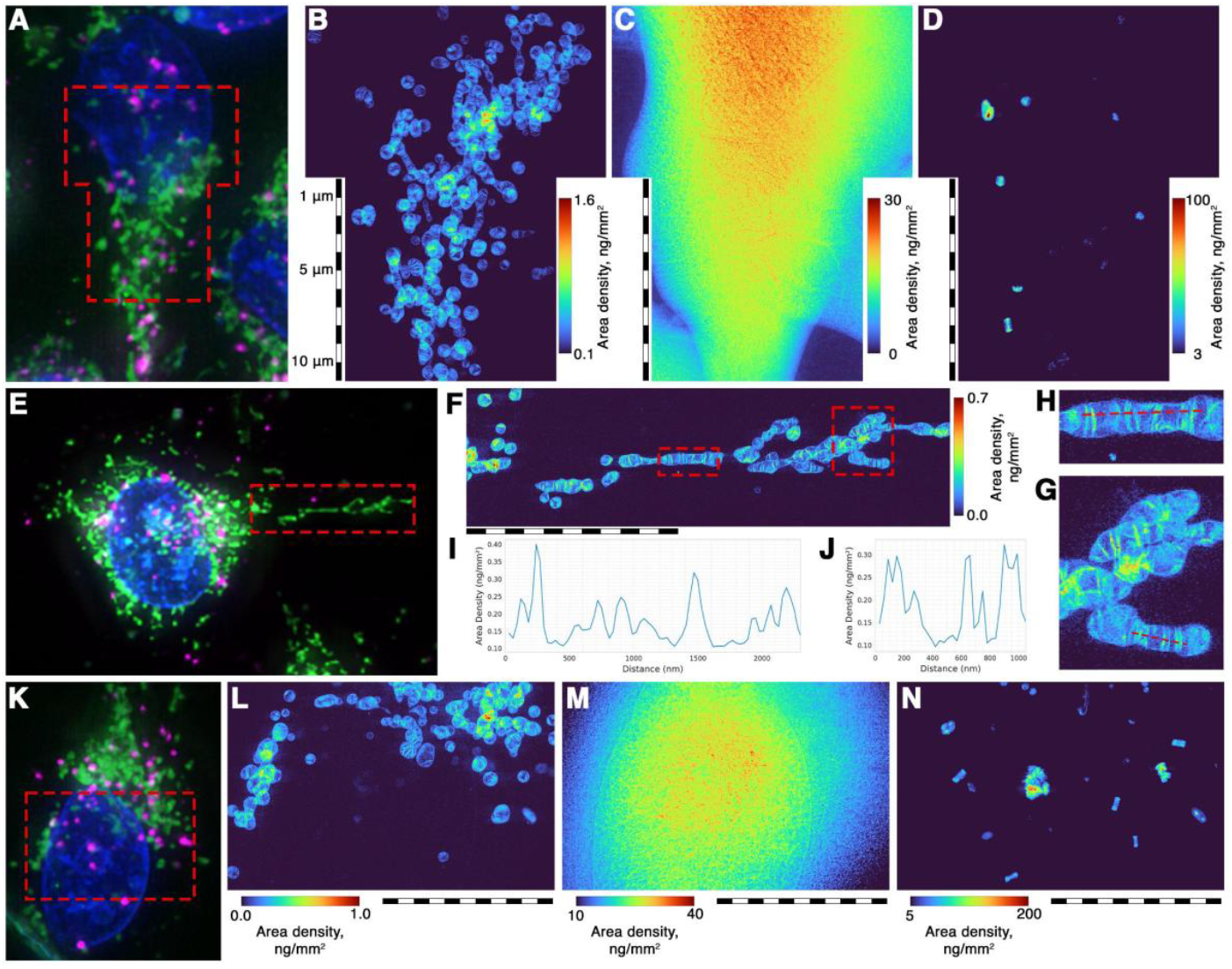
Elemental cryo-XRF-N and optical cryo-FLM correlative images of MDA-MD-231 breast cancer cells exposed to PEG-GdTbF_3_ nanohybrids and Ir-Mito mitochondrial luminescent probe. Navigation through the acquired cryo-FLM MIP images allow to choose cryo-XRF-N acquisition on the very same location as shown in **(A-D)** over nearly the entire cell surface scanned with a 50 nm X-ray pixel size, projected concentration as areal mass is given in ng/mm^2^. It shows clear visual correspondence between the mitochondrial luminescent signal (**panel A and K**, green channel) and the iridium XRF signal in **panel B** and **L** respectively, with some internal features that are discernable in mitochondria. Cluster of GdTbF3 nanoparticles are also registered for example Gd (**panel D and N**) but Tb distribution is strictly similar to Gd and the Gd/Tb ratio inside the cell compartments is found close to the native value of the synthesized nanohybrids. The potassium (K) signal depicts well the shape of the whole cell profile and thickness with a quite homogeneous signal (**panel C and M**), indicative of a very good chemical preservation of the ionic content and of the membranes. When vitrification is bad or when osmotic damages from the rinsing buffer occurs generally the cell K and other labile ions induce leaching of ionic content and resulting in a “holey structure” of the K map. Detailed region of interest at organelle level can be targeted using the cryo-FLM MIP images **(E)** and higher XRF scans (30 nm pixel size) can be performed (**panel F, G and H**). This allows to more clearly depicts lamellar structure inside mitochondria with structures and spacing dimension (**panel F-I**) fully compatible with what is observed in optical fluorescence nanoscopy i.e. cristae which has never been reported so far in XRF nanoimaging. Further the Gd or Tb XRF maps simultaneously registered gave an exact corresponding localization with the lysosome found in this field-of view (data not shown).

**Figure 3:**
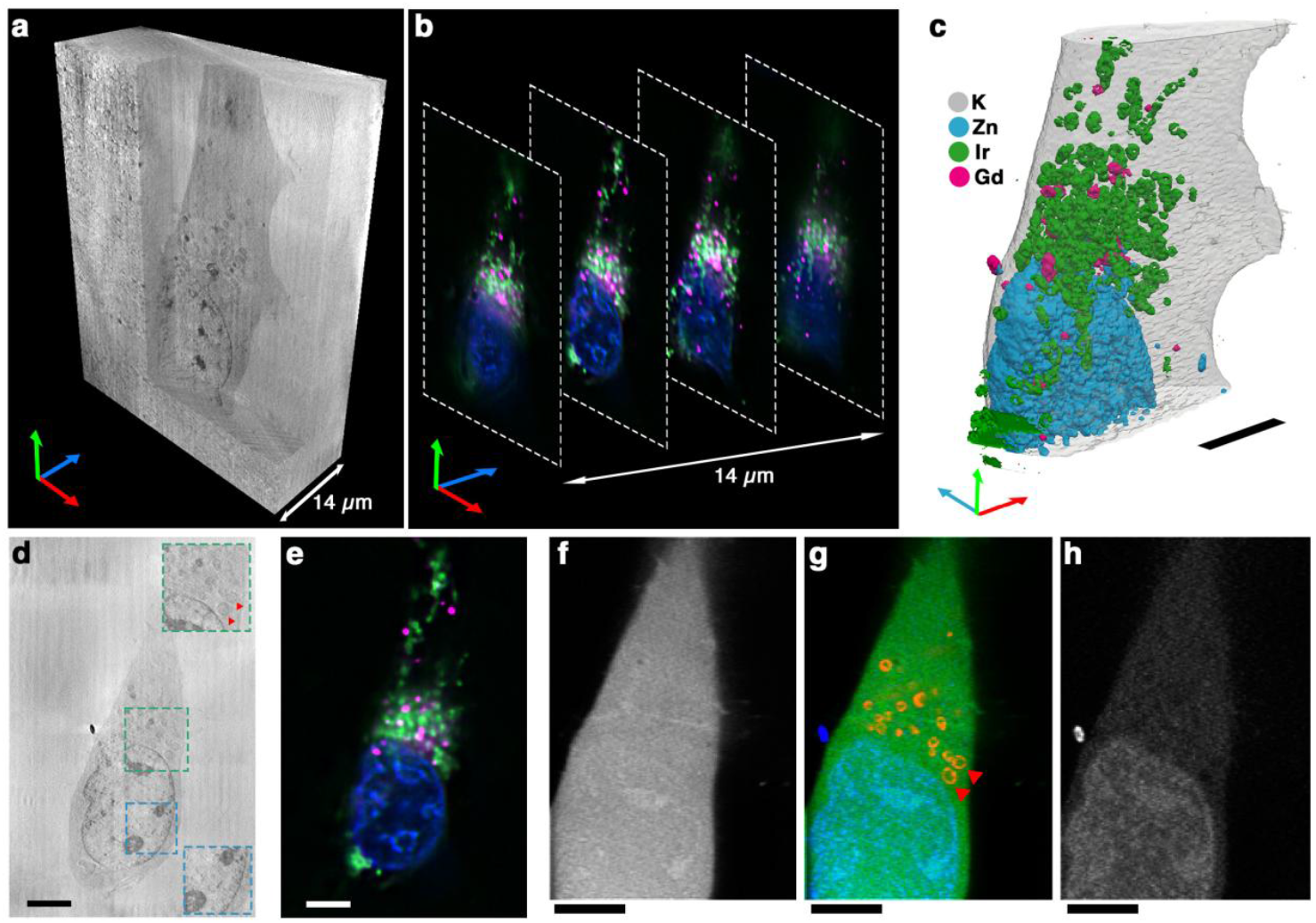
Three-dimensional visualization of the cellular structures with the proposed cryo-correlative workflow. **(a)** volume rending of the phase contrast nano tomography data.**(b)** slices through the cryo-optical fluorescence microscopy data. **(c)** volume rendering of the elemental distribution obtained with X-ray fluorescence nano tomography. **(d)** central slice through the volume of the phase contrast data. The red triangles indicate the iridium compound. **(e)** slice of cryo-optical fluorescence microscopy corresponding to the slice in *d*. **(f)** slice through X-ray fluorescence tomogram of a K elemental line. **(g)** color composite of the slice through X-ray fluorescence tomogram with green color used for the K elemental line, blue color for the Zn elemental like, and red color for the iridium elemental line. **(h)** slice through X-ray fluorescence tomogram of a Zn elemental line. Scale bar 10 µm

The proposed workflow cannot yet identify the molecular target of Ir-Mito. However, Sorvina et al. suggested a possible mechanism through a protein or a lipid association [11], that can now be explored through the present correlative nanoimaging workflow. An asset of XRF is its quantitative assessment and it can be noted that the areal mass of iridium in the inner mitochondrial structures remains in the same range (0.1-0.3 ng/mm^2^) in different elemental maps acquired (Fig. 2, Supplementary Fig. 3 & 4). For example, approximately 9000 iridium atoms were detected within individual lamellar structures only a few tens of nanometers in size, corresponding to mitochondrial cristae regions (Fig. 2, Supplementary Fig. 3 & 4). Similarly, the concentration of Gd and Tb, which constitute the core of the nanohybrids, is also of interest for further in-depth studies on their intracellular uptake and, in particular, their sequestration into lysosomes. For example, up to 15 fg of Gd and 2 fg of Tb can be found presently in lysosomes, the Tb/Gd remaining close to the 0.1% of the raw nanohybrids. Assuming each nanohybrid of 10 nm in diameter [12] contains approximately 10^6^ atoms of Gd, we estimate that up to 60 nanohybrids are stored in lysosome of MDA-MB-231 cells after 24 hours of exposure under our experimental conditions.

The 3D elemental nanoimaging performed on vitrified MDA-MB-231 cells allowed for the depiction of the intracellular mitochondrial network through the iridium signal matching the optical fluorescence signal of the cryo-FLM and it confirms that within the cellular volume detected Gd and Tb atoms of nanoparticles are confined mostly into lysosomes (Fig. 3). A compromise was made to ensure a reasonable analytical time of approximately 20 hours while prioritizing a higher number of projections (81 in our case). This approach led to tomographic acquisition being performed with a step size of 130 nm. The three volumes can be co-registered to offer a quantitative elemental distribution across the cell, aligned with structural details observed either through X-ray phase contrast nanoimaging—highlighting nuclear and nucleolar regions while beginning to reveal certain cytosolic intracellular structures—or through cryo-FLM, where live labeling of cellular organelles is performed prior to vitrification (Fig. 3).

The proposed integrated method enables us to both image the labeled subcellular compartments and organelles, alongside sub-organelle level volumetric mapping of elemental composition within entire frozen-hydrated cells. This minimally invasive approach highlights the preservation of the ultrastructure and chemical integrity of the native cellular interior while delivering detailed structural and chemical insights. The current spatial resolution of 130 nm remains a limitation, but it is reasonable to anticipate that achieving full 3D X-ray fluorescence nanotomography of an entire adherent cell, vitrified in its near-native state, with a spatial resolution of 50 nm will soon be possible. This advancement could be realized within just one or two synchrotron shifts (equivalent to 8–16 hours).

Beyond the recent advent of high-throughput techniques for single-cell studies in the “omics” field, our new implementation demonstrated cryo-correlative X-ray nanoimaging opening possibilities to explore quantitatively the volumetric elemental distribution at organelle level on an entire frozen-hydrated cell and expand the knowledge on the chemical landscape of the interior of a cell. As a source of inspiration, workflow powerfully developed in the cryo-electron microscopy (EM) domain to merge ultrastructural changes/information to intracellular events could be fully compatible through further focused-ion beam milling and cryo-electron tomography at nm level. Our approach unlocks the untapped potential of developing a platform of molecular probes that are differentiated by the presence of heavy elements and tailored to target specific organelles. This combinatorial approach offers a suite of molecular probes designed to achieve potent multiplexing imaging, that provides both detailed structural insights and quantitative information. An excellent review, which the reader is referred to, spans recent studies where optical fluorescence microscopy and X-ray fluorescence were combined to study cellular systems [18]. Serpell et al. show that carbon nanotubes can serve as flexible “containers” for xenobiotic elements, enabling multichannel XRF imaging [19]. However, their demonstration was limited to chemically fixed cells at a spatial resolution of about 2 µm and lacked both organelle-specific targeting and optical emission properties. To our knowledge, synchrotron radiation X-ray fluorescence nanoimaging (SR-XRF-N) with elemental multiplexing in vitrified cells using organelle-specific, multi-element luminescent organometallic probes has not yet been achieved. Such an approach would provide a direct “elemental paint” for organelles whose native elemental composition is largely indistinguishable from the surrounding cytoplasm, thereby generating an enhanced elemental internal contrast required for their unequivocal identification. As a proof of concept, we have selected three additional species, alongside Ir-Mito, as a small family of luminescent molecular probes with defined cellular targets and differentiated by the presence of various heavy elements that can be useful XRF reporters (Supplementary Fig 2 & Supplementary Fig. 5). The complex Re-ER bears the heavy element rhenium and was previously demonstrated to accumulate within the endoplasmic reticulum [20]. On the other hand, lysoBODIPY-Br and lipoBODIPY-I are based around the fluorescent 4,4-difluoro-4-bora-3a,4a-diaza-*s*-indacene core and feature a bromine and iodine atom, respectively. Furthermore, lysoBODIPY-Br is functionalised with a morpholine-appended side chain for lysosome targeting, whereas lipoBODIPY-I has demonstrated affinity for lipophilic regions in brain tissue [21]. In this first proof-of-concept, a key limitation is that the emission profiles of these probes have significant spectral overlap in the visible region making them indistinguishable by cryo-FLM. Therefore, they cannot yet serve as true bimodal agents for combined cryo-FLM and SR-XRF-N. However, as previously mentioned, our current aim is to demonstrate that these probes deliver strong intracellular elemental contrast of labelled organelles, allowing their unambiguous detection in XRF maps. The characteristic X-ray lines selected for imaging with iodine Lβ at 4.22 keV, iridium Lβ_1_ at 10.71 keV, and bromine Kα at 11.9 keV fall outside the emission ranges of the major endogenous elements, so they generate virtually no spectral overlap in XRF analyses. SR-XRF-N with a 17 keV excitation beam confirmed this advantage: in HeLa cells, lipoBODIPY-I accumulated in sharply defined spherical inclusions, while the lysoBODIPY-Br localized into a separate set of spheres corresponding to lysosomes (Fig. 4a). The same cells were also incubated with the Re-ER complex. Residual rhenium remains detectable inside the cells, with slightly higher concentrations in the perinuclear region, consistent with the initial labelling of the endoplasmic reticulum (Fig. 4a). We have already shown that this probe enters cells by passive diffusion and binds weakly to its target, as indicated by the rapid loss of luminescence signal upon washing [20]. The XRF pattern is therefore expected: after incubation, the cells were rinsed with PBS, transferred to dye-free culture medium, and, immediately before vitrification, briefly washed in trace-element-free ammonium acetate buffer (290 mOsmol/L, pH 7.4). Therefore, most of the Re-ER weakly bound to the endoplasmic reticulum was washed away. Cells were in all cases pre-labelled with Hoechst 33342 before vitrification, enabling SR-XRF-N imaging and we could unequivocally identify a mitotic cell in which the chromosomes are organized into a metaphase plate (Hoechst channel not shown). The phosphorus map in such mitotic cells mirrors this arrangement, tracing the chromosome band aligned across the spindle mid-zone (Fig. 4b). As expected, zinc is uniformly distributed throughout the cytoplasm, reflecting the breakdown of the nuclear envelope during metaphase. Concurrent XRF acquisition with our set of probes provides intrinsic elemental contrast that delineates mitochondria, lysosomes and lipid droplets distribution within mitotic cells (Fig. 4b). Extensive mitochondrial remodeling is a hallmark of mitosis, becoming most pronounced at metaphase through a spatial shuffling of mitochondria within the mother cell where the network fragments and relocates toward the cell cortex. The iridium-labeled SR-XRF images in our study (Fig. 4b) faithfully reproduce this redistribution, corroborating the well-documented behaviour of mitochondria during cell division.

**Figure 4:**
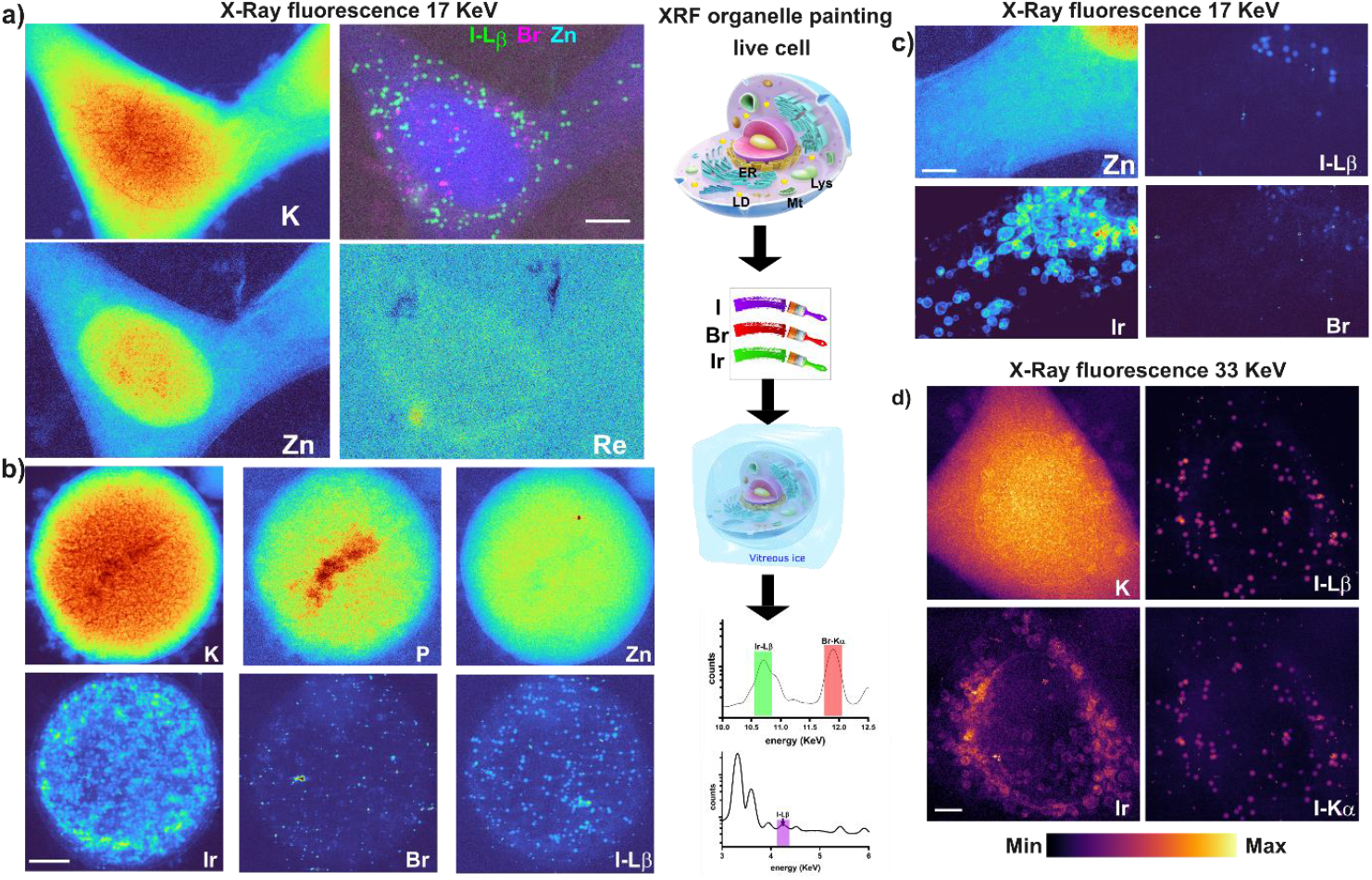
Organelle elemental multiplexing cryo-XRF-N images of HeLa cancer cells exposed Ir-Mito, Re-ER, lysoBODIPY-Br and lipoBODIPY-I. All cells were incubated with Hoechst immediately before vitrification (signal not shown). **(a)** Frozen-hydrated, proliferating HeLa cell simultaneously labeled, prior to vitrification, with Re-ER, lysoBODIPY-Br and lipoBODIPY-I. X-ray fluorescence (XRF) map acquired with 50 nm pixels and a 30 ms dwell time. **(b)** Frozen-hydrated mitotic HeLa cell exposed simultaneously prior vitrification to Ir-Mito, lysoBODIPY-Br and lipoBODIPY-I. Acquisition conditions identical to (a). **(c)** SR-XRF-N (30 nm spatial resolution) of a frozen-hydrated cytoplasmic extension from a HeLa cell treated with the same probes as in (b) prior to vitrification. **(d)** The ID16A nano-imaging beamline can also operate at 33.6 keV with a 30 nm focal spot, matching the 17 keV focus used above. This higher excitation energy enables detection of the iodine K-emission line while retaining sensitivity to iodine and iridium L-lines and to endogenous cellular elements.

Further, high-resolution (30 nm) scanning of a cytoplasmic extension of an HeLa cell shows that the use of the organometallic probes for XRF multiplexing of various organelles does not impact, for example, the localization of Ir-Mito inside mitochondria, as revealed above in the present work, and that they can be mixed without interference between each other and between physiological cellular elements (Fig. 4c). Finally, using the high energy capabilities of the ID16A beamline providing a 30nm focus not only at 17 keV but also at 33.6 keV above the K-edge of iodine, the X-ray fluorescence K*α* emission line at 28.6 keV is well detected concomitantly with the iodine Lβ at 4.22 keV sharing as expected the very same distribution in lipid droplets (Fig. 4d). Although this high energy excitation is not optimal for elements detected at much lower energy like for potassium or even zinc or the L-lines of iridium, still some decent contrast and definition is observed when compared to 17 keV excitation. These proof-of-concept pave the way for engineering an expanded palette of probes by combining a variety of organelle-targeting functionalities with a chemical structure containing heavy atoms amenable for XRF detection. This approach further enhances the multiplexing capacity of synchrotron XRF “painting” for the simultaneous visualization of multiple sub-cellular compartments.

Overall, the present workflow, with its capacity for continuous improvement in acquisition speed, atomic sensitivity, and ultimate X-ray spot resolution, paves the way for new possibilities in 2D/3D chemical information within the intracellular environment. Notably, it facilitates the incorporation of heavy elements, such as halogens and *d-* or *f*-block elements, within the chemical design of the probes to be developed as reporters in SR-XRF-N, as a step toward a complementary structural approach for the quantitative investigation of specific intracellular proteins, metabolites, and pathways.

## Author contributions

D.K., F.L, P.C. and S.B. conceived and designed the study. D.K., R.S., P.C. and S.B. performed the experiments. M.H., M.M. E.D., B.S. and S.P. designed and synthesised the molecular probes. D.K., L.C., R.S., C.G., S.P., M.M., F.L., E.D., B.S., M.H., P.C. and S.B. analyzed the data. D.K., S.P., P.C. and S.B. wrote the paper. Funding acquisition: F.L. and S.B. All authors discussed the results, assisted in the preparation, and contributed to the manuscript. All authors approved the final version of the document.

## Methods

### 1 Naohybrids synthesis and characterization

For synthesis, 3.64g of GdCl_3_, 6 H_2_O (9.8 mmol) and 0.41 g of TbCl_3_, 6 H_2_O (1.1 mmol) are dissolved in 5g of ethylene glycol. In the meantime, 875 µL (25 mmol) of an aqueous solution of HF (50 % wt) are added to 20 g of pyrrolidinone. The HF solution is then added to the lanthanide salts mixture. The mixture is heated under pressure in the autoclave at 170°C during 1h30. NPs are precipitated in acetone and washed in methanol by centrifugation-redispersion cycles. Then, they are dispersed in water. 50g of an aqueous suspension (10 % wt) of Gd_0.90_Tb_0.10_F_3_ NPs (2.3.10^−2^ mol of Ln) are sonicated in an ultrasonic bath. This suspension is added slowly to a solution of OMe-PEG phosphonic acid (7.7g (7.7.10^−3^ mol) in 85 mL of water). The mixture is heated at 80°C during 24h and purified by dialysis. After purification the suspension is freeze-dried and NPs kept as a powder. NPs were characterized by X-ray powder diffraction, High-resolution transmission electron microscopy and dynamic light scattering. They crystallize in a GdF_3_ orthorhombic structure and have an inorganic core with an ellipsoidal shape with an average size of 10 nm. The hydrodynamic diameter of PEG-modified NPs is evaluated at 24 nm. The presence of the ligand PEG is characterized by Fourier Transform Infrared Spectroscopy and Thermogravimetric analysis. The number of ligands was estimated to be 2700 ligands per NPs.

#### Molecular probes synthesis and characterisation

Ir-Mito [22], Re-ER [20] (also commercially available from Biosensis as LipoFluor-MR™ and LipoFluor-ER™, respectively) and lipoBODIPY-I were prepared according to previously published procedures [23]. lysoBODIPY-Br was prepared according to the synthetic scheme detailed in Supplementary Fig. 5.

### 2 Cell Culture

The MDA-MB-231 human breast adenocarcinoma cell line, Human cervical cancer HeLa cells were purchased from the American Type Culture Collection (ATCC, Rockville, MD), and were maintained in monolayer culture in DMEM medium with Phenol Red/Glutamax I (Gibco) supplemented with 10% fetal calf serum (Gibco) in a 75 cm2 flask at 37 °C under 5% CO2/air-humidified incubator. Cells were plated onto silicon nitride windows (5×5 mm wide, 200 μm thick silicon frames with 1.5×1.5 mm wide and 500 nm thick silicon nitride membranes coated with graphene that has been shown to provide in cryo-electron microscopy a high thermal conductivity promoting heat dissipation of the ice with concomitant improved thermal stability under electron irradiation at cryogenic temperature [24,25]. Although the graphene coating already improves hydrophilicity of the Si_3_N_4_ membrane, the cellular adhesion and growth is by far enhanced through an air plasma treatment using a Henniker HPT-100 plasma machine (50% power, 20 seconds, Henniker Plasma, Runcorn, UK) resulting in a hydrophilized membrane. This process avoids the classical requirement of a coating substrates, such as Poly-L-Lysine, Fibronectin or collagen for cell adhesion. The membrane has an additional small square orientation Si_3_N_4_ membrane in one of the corner of the Si frame (Silson Ltd, UK) and are deposited in each well of a 4-well plate chamber [26]. The cells (∼3 × 10^3^ cells deposited onto the membrane) were allowed to attach and grow overnight in a regular medium [26]. The following day, cells were incubated for 24h with 670 µM of nanohybrid formulation described above.

### 3 Sample preparation

After the 24h of incubation with nanohybrid, cells were labelled with vital fluorescent markers, Hoechst 33342 NucBlue® Live ReadyProbes® Reagent (Thermo Fisher Scientific, MA, USA) was used to label cell nuclei and LysoTracker® Deep Red (Thermo Fisher Scientific, MA, USA) at 50 nM was used to label acidic organelles (specifically lysosomes), following the manufacturer’s instructions. Ir-Mito (20 µM) was used to label mitochondria, Re-ER (50 µM) was used to label endoplasmic reticulum, lysoBODIPY-Br (20 µM) was used to label lysosomes, and lipoBODIPY-I (20 µM) was used to label lipid droplets, all diluted in cell culture media from DMSO stock solutions and used with an incubation time of 30 min at 37 °C to provide simultaneous labelling where indicated. Finally, the membrane where seated live cells were quickly rinsed in a trace-element-free ammonium acetate buffer (290 mOsmol/L, pH 7.4), blotted with no. 1 Whatman filter paper prior to freezing and then automatically plunged into liquid ethane (Leica EM GP) before being transferred to liquid nitrogen storage [26]. The silicon nitride membranes were stored in liquid nitrogen using a custom-designed 3D-printed puck system with nine slots. Each slot housed a 3D-printed storage box capable of holding three silicon nitride membranes. These storage boxes were designed to match the standard format of routine storage of pucks for cryo-EM grid boxes (Supplementary Fig. 1). The 3D-printed storage elements are available from the authors on reasonable request. Storage of the frozen hydrated samples in liquid nitrogen or at cryogenic temperatures on the cryo-fluorescence light microscope or synchrotron X-ray fluorescence cryo-nanoimaging setup preserve cell from ice crystal formation/contamination. It sometimes remains non-trivial with Si_3_N_4_ membrane to optimize ice thickness, and, cellular distribution that should not be too dense to avoid shadowing and overlapping signal from neighboring cells at high tilt angles when tomography is considered as in the present work. In our cases, ice thickness achieved is within the 10 µm range average from X-ray transmission intensity measurements on the 1.5mm square area of the membrane and an initial seeding density of 3.10^3^ cells was found appropriate (see above). All further manipulation required to maintain the sample well below the devitrification temperature i.e. below 130 K [27].

### 4 Cryo-Fluorescence Light Microscope

Prior to be transferred for synchrotron X-ray fluorescence cryo-nanoimaging, plunge-frozen cells on the Si_3_N_4_ membranes, were imaged using widefield cryo-fluorescence light microscope (cryo-FLM) Leica cryo-CLEM Thunder® system equipped with a ceramic-tipped, 0.9 NA, 50 X lens. The brightfield and band pass filter cubes of GFP (emission, l = 525 nm), DAPI (emission, l = 477 nm), Texas Red (emission, l = 619 nm) and Y5 (emission, l = 660 nm) were used. A complete mosaic of the 1.5 × 1.5 mm Si3N4 active area containing vitrified cellular region of interest was registered with the collection for each field of view as a Z-stack projections over ∼ 10 µm for each channel, Nyquist sampling was used to set the z-interval between every slice and Small Volume Computational Clearance (SVCC) from Leica LAS X THUNDER package was applied for fluorescent image deconvolution and blurring reduction on the cryo-FLM image stacks. In addition to thunder processing, maximum projection was applied using Las-X software (Leica Microsystems).

### 5 Sample handling and measurement conditions at the synchrotron experiments

Synchrotron experiments were performed on the ID16A Nano-Imaging beamline at the European Synchrotron Radiation Facility (Grenoble, France) [28]. The beamline is optimized for cryogenic X-ray fluorescence imaging as well as coherent hard X-ray imaging (in-line X-ray holography and X-ray ptychography). Featuring two pairs of multilayer coated Kirkpatrick-Baez (KB) focusing mirrors, the beamline provides a pink nanoprobe (ΔE/E≈1%) at two discrete energies: E = 17 keV and 33.6 keV. The capability of the beamline to operate under cryogenic conditions makes it particularly useful for bioimaging applications. The cryogenic experiments are performed at approximately 120K.

Manipulation of our homemade cryo-puck and cryo-boxes can be manipulated as regularly done in cryo-EM or X-ray crystallography labs. Pucks are manipulated with specialized tongs with hemostat-style closure and transferred into dedicated LN2 container for handling the Si_3_N_4_ boxes. Then, silicon nitride membranes were moved to a sample cryo-manipulation system (EM VCM, Leica Microsystems, Germany), where they remained submerged in liquid nitrogen at all time. Next, each membrane was mounted and clamped vertically within a dedicated aluminum-based sample holder in liquid nitrogen. The entire assembly was subsequently transferred to the docking port of the high-vacuum ID16A endstation using a vacuum cryo-transfer system (EM VCT500, Leica Microsystems, Germany), which maintained the samples under vacuum in a cryo-shielded environment at T = 110 K.

Following this, the membranes holding the plunge-frozen hydrated cells were mounted under high vacuum (10^−8^ mbar) onto a piezo nano-positioning stage with six short-range actuators, regulated via the metrology of twelve capacitive sensors at all steps. Throughout this procedure, the sample temperature was kept well below the devitrification threshold for biological samples (approximately 130 K).

### 6 Synchrotron X-ray fluorescence cryo-nanoimaging

We used 17 keV excitation energy for most measurements but also the 33.6 keV excitation was used for some measurements in the multiplexed experiments. In our experiment we have used the X-rays focused down to 30 nm spot size providing a flux of 2 × 10^11^ ph/s [28]. The spot size was determined with a lithographic sample consisting of a 10 nm thick, 20 × 20 µm^2^ square of nickel deposited on a 500 nm thick silicon nitride membrane. This test target was rapidly scanned along orthogonal directions and the spot size determined using the knife-edge test. An ultra-long working distance optical microscope was used to bring the sample to the focal plane (depth-of-focus ± 3 µm) and to position the cryo-FLM regions of interest in the X-ray beam. The samples were scanned with an isotropic pixel size of 50 and 30 nm with 50 ms dwell-time, unless indicated otherwise. The X-ray fluorescence signal was detected with two custom energy-dispersive detectors, a ME-7 multi-element SDD (Hitachi Ltd.) and an ARDESIA-16 spectrometer based on monolithic SDD array [10] were used. Detectors are positioned at either side of the sample, holding a total of 23 silicon drift diodes, coupled with data stream is handled with an XIA FalconX processor for (XIA LLC, Hayward, CA, USA) resulting in a solid angle of 1.2 steradian. Each detector channel of the FalconX list-mode data stream is written to disk separately, and then summed in a pre-processing step before analysis.

The quantitative data treatment of the SXRF data was performed with in-house Python scripts using the PyMCA library [29] using the fundamental parameter method with the equivalent detector surface and geometry determined by calibration with a thin film reference sample (AXO DRESDEN GmbH) and considering a ice matrix of thickness 10 µm. The resulting elemental areal mass density maps (units: ng.mm^−2^) were visualized with ImageJ (National Institutes of Health, Bethesda, Maryland, USA).

For 3D XRF imaging, we have obtained elemental maps at 81 different tomographic angles, mapping the region of interest of 22 × 28 μm^2^ with a 130 nm step size and 25 ms dwell time per point. A 40° angular range was avoided resulting in a missing wedge that is common for this type of samples. A single fluorescence map took ca. 15 minutes and a full tomographic scan took ca. 20 hours. Obtained 2D fluorescence maps were carefully aligned and processed into 3D volumes using a Maximum-Likelihood Expectation-Maximization (MLEM) [30] algorithm implemented with ESRF inhouse software.

### 7 Synchrotron X-ray phase contrast imaging

The X-ray phase contrast imaging (PCI) was done with in-line holography [31]. The sample was positioned downstream of the focus to achieve 25 nm pixel size. The X-ray holograms were recorded with indirect imaging system based a XIMEA sCMOS detector with 6144×6144 pixels. The images were binned 3-by-3 resulting in 51.2 µm by 51.2 µm field of view. X-ray holograms were collected in the Fresnel region at four different focus-to-sample distances to assure efficient phase retrieval. Phase retrieval was performed using the contrast transfer function approach, implemented in ESRF inhouse code using the GNU Octave language. This approach is well adopted to weak phase objects such as biological cells embedded in ice. Obtained 2D phase maps acquired at different tomographic angles were processed into a 3D volume using Nabu software package developed at the ESRF.

### 8 Synchrotron X-ray imaging dose estimation

Assuming vitreous ice with a density of 0.94 g/cm3 and membrane silicon nitride with an approximate composition SiN and a density of 3 g/cm3 we can estimate the deposited dose during the X-ray imaging experiment. For the holographic tomography at the 25 nm pixel size the cell has received 2 MGy at 25 nm. For the X-ray fluorescence tomography with the 130 nm pixel size the cell has received 40 GGy, and for the high resolution 2D X-ray fluorescence scan at 30 nm pixel size the cell has received 3 GGy.

### 9 Correlative analysis

To analyse the data we first performed data alignment using eC-CLEM plugin-in from the Icy software platform version 2.5.2.0 [32]. We used cryo-light fluorescence and XRF projection images correlating features such as the nuclear shape well depicted in optical fluorescence through the Hoechst live staining and in the XRF Zn map that display also clearly shape and contour of the cell nucleus. The 3D registration between the X-phase contrast holotomography and the wide-field cryo-FLM 3D z-stack of the very same single cell were also registered using Icy software and eC-CLEM plugin using nuclear feature visible in the X-ray phase contrast holotomography and the Hoechst staining from the 3D z-stack acquired data.

