## Supplementary Information for "Cryo-Correlative Light and X-ray microscopies: Expanding the Intracellular Chemical Map"

1 European Synchrotron Radiation Facility, F38043, Grenoble Cedex 9, France.

2 Université de Lyon, École Normale Supérieure de Lyon, CNRS UMR 5182, Université Claude Bernard Lyon 1, Laboratoire de Chimie, Lyon F69342, France

3 Clinical and Health Science, University of South Australia, Adelaide, 5000, SA, Australia

4 School of Molecular and Life Sciences, Curtin University, Perth, Western Australia 6102, Australia.

5 Université Grenoble Alpes, Synchrotron Radiation for Biomedical Research (STROBE), Inserm UA 7, 71 Avenue des Martyrs, Grenoble Cedex 9 38043, France.

### Supplementary Materials and Figures

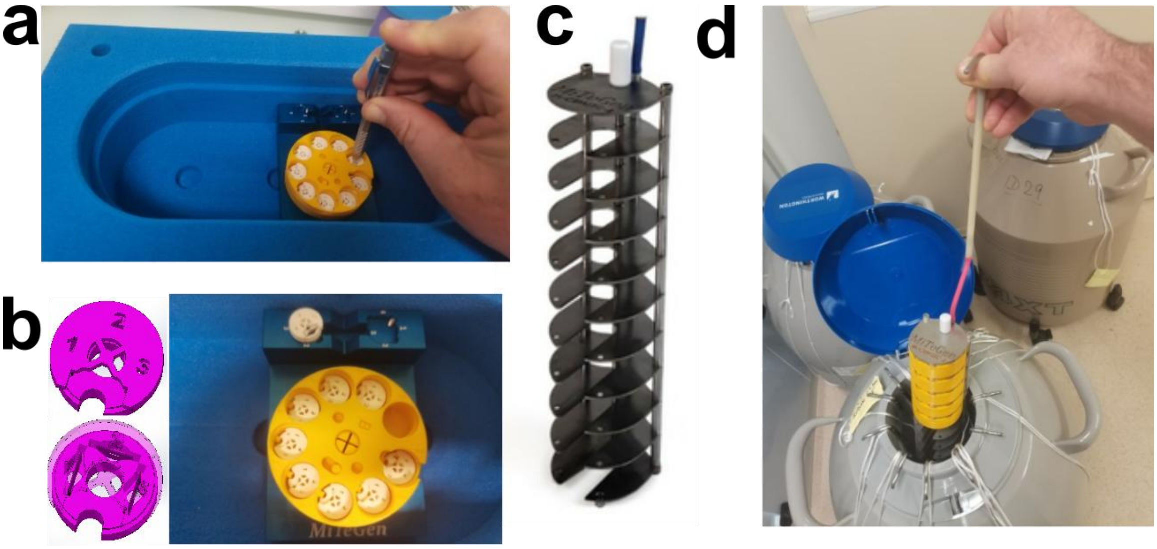


**Fig. S1 | The cryo-storage system designed and developed for the preservation of cryo-SR-XRF-N samples**. **(a)** The cryo-storage system consists of 3D-printed puck that can be manipulated within a commercially available foam dewar and cryo-EM puck holding metallic insert for easy sample transfer. **(b)** The storage pucks are designed to hold nine 3D-printed cryo-box, each having 3 slot positions with identification numbers for the storage of vitrified samples on $Si_3N_4$ membranes in liquid nitrogen storage dewars. The lid is rotated with standard laboratory tweezers or a Criterium mechanical pencil. The cryo-box is of standard size for all common Cryo-EM sample mounting such as in automatic plunge freezer machine, and storage devices. **(c & d)** The 3D-printed pucks are vertically stacked in a nine-shelved storage canister for cryo-EM grid-box storage pucks and transfer in long term liquid nitrogen storage dewars.

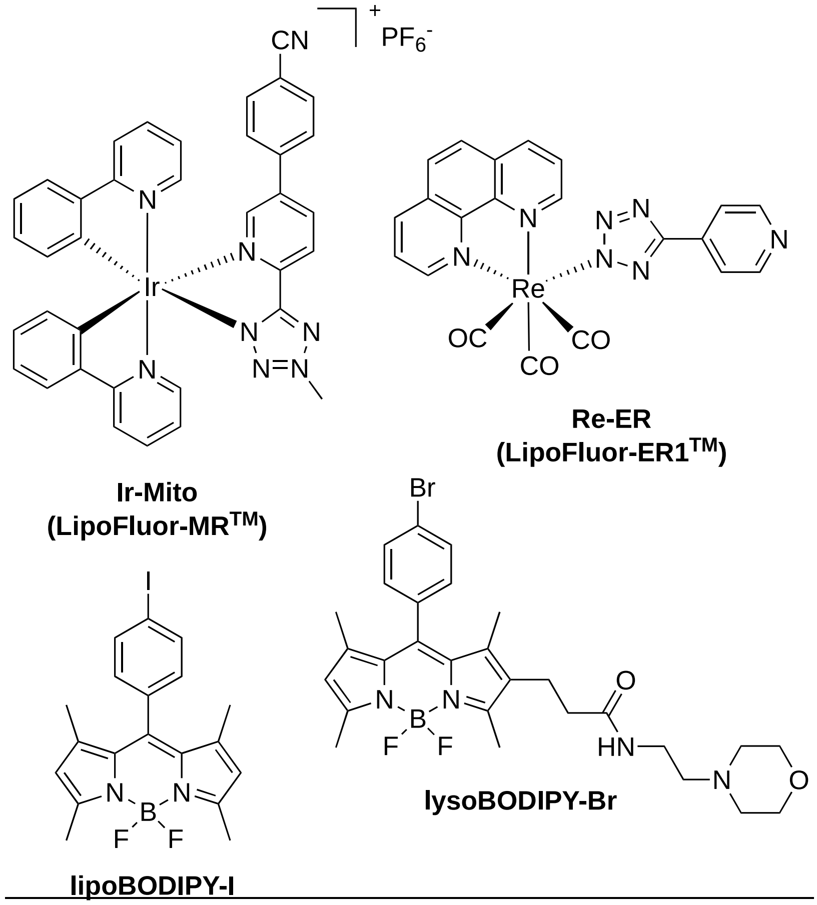


**Fig. S2: Structures of the luminescent probes used in this work**.

Ir-Mito is an iridium complex targeting mitochondria. Re-ER is a rhenium complex targeting the endoplasmic reticulum. lysoBODIPY-Br and lipoBODIPY-I are compounds that preferentially accumulate within lysosomes and lipid droplets, respectively.

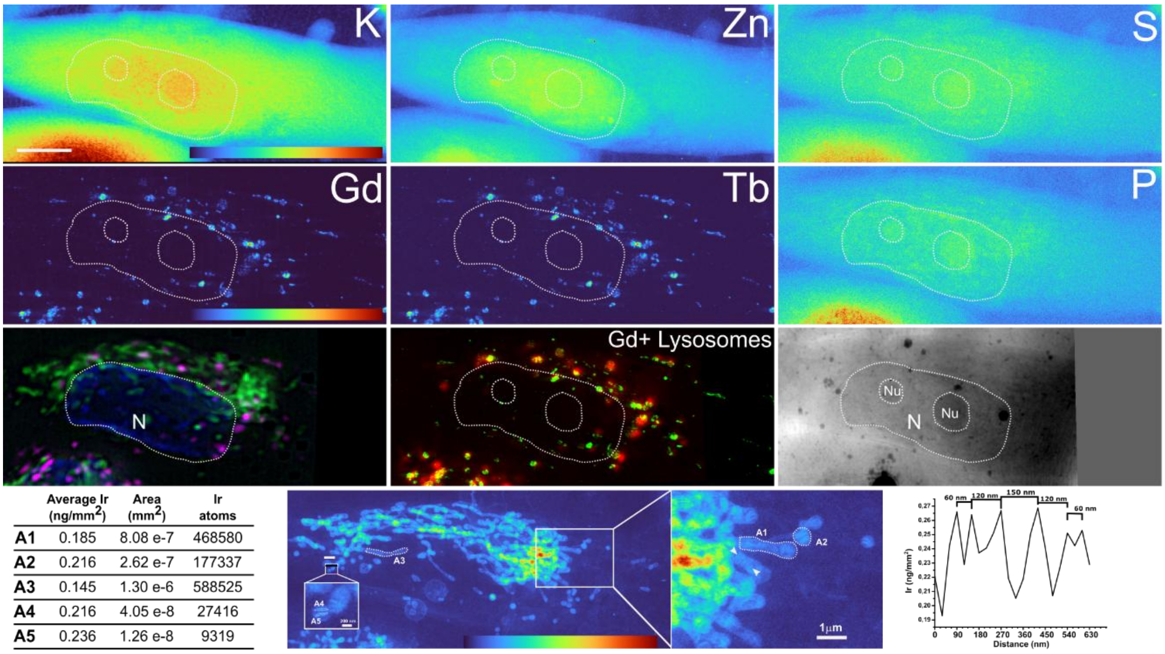


| | Average Ir (ng/mm²) | Area (mm²) | Ir atoms |
|---|---|---|---|
| A1 | 0.185 | 8.08 e-7 | 468580 |
| A2 | 0.216 | 2.62 e-7 | 177337 |
| A3 | 0.145 | 1.30 e-6 | 588525 |
| A4 | 0.216 | 4.05 e-8 | 27416 |
| A5 | 0.236 | 1.26 e-8 | 9319 |

**Fig. S3: High-resolution cryo-XRF-N images (50 nm pixel size) of entire cell with correlated registration between cryo-FLM, SR-XRF-N and X-ray phase contrast nanoimaging following the proposed cryogenic workflow.**

The cells imaged were MDA-MB-231 triple-negative aggressive breast cancer cells, exposed to $GdTbF_3$-PEG nanohybrids[11,13] and fluorescent live labelling of the cellular organelles. Elemental distribution within single whole breast cancer cells in close contact to neighbor one is displayed for potassium **(a)**, zinc **(b)**, sulfur **(c)**, gadolinium **(d)**, iridium **(e)** and phosphorus **(f)** with their respective projected areal mass concentration (ng/mm$^2$). Region of interests from the iridium map and respective zoomed area are displayed **(g-j)** that allow to distinguish some lamellar structured into mitochondria with spacing in the 60-120 nm distance compatible with cristae structures, and last but not least possibilities to derived number of iridium atoms within various regions (A1-A5 areas) of mitochondria (left side table). The corresponding cryo-FLM using computational clearing processing and maximum integrated intensity of image stack (MIP) shows cell nucleus in blue, mitochondria in green and lysosomes in pink **(k).** Registration and merging of the Gd XRF map and the cryo-FLM pink channel (lysosomes) allow to observe some co-localization **(l)** while hard X-ray phase contrast shows cellular and nuclear contrast and shape despite extreme low absorption of the cell at 17 keV **(m)**. Scale bar 5 µm.

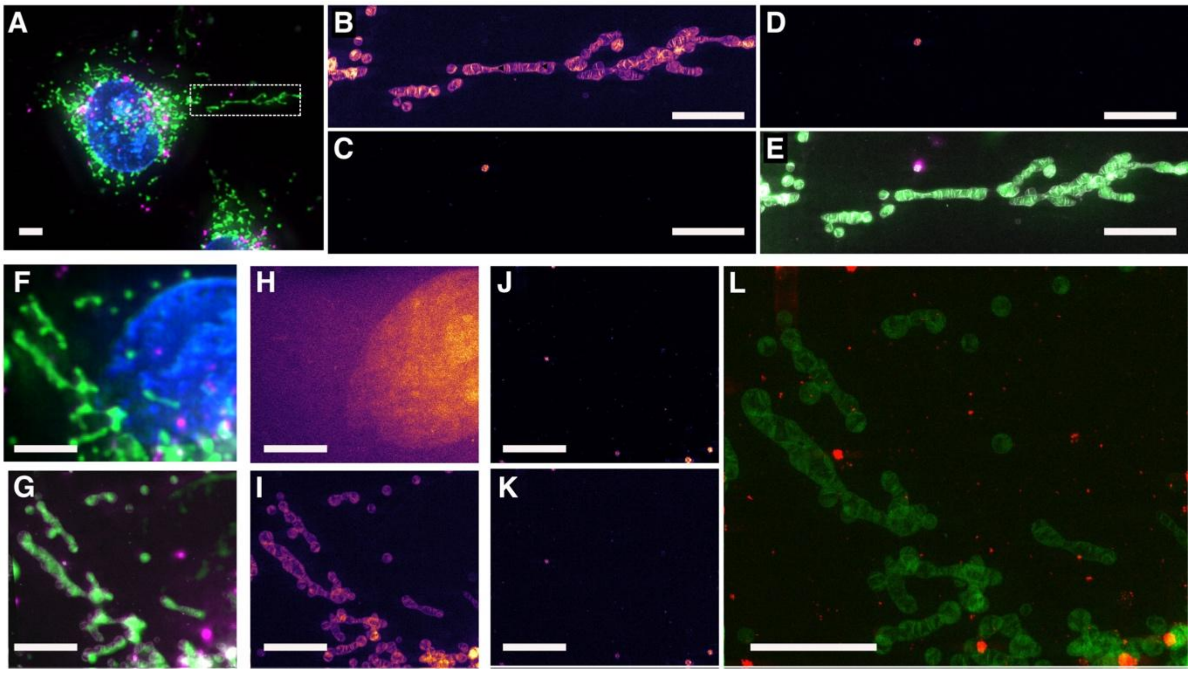


**Fig. S4: High-resolution (30 nm pixel size) cryo-XRF-N images of entire MDA-MB-231 breast cancer cell with correlated registration between cryo-FLM and SR-XRF-N following the proposed cryogenic workflow. Scale bars 5 µm.**

The cells imaged were triple-negative aggressive breast cancer cells, exposed to $GdTbF_3$-PEG nanohybrids[11,13] and fluorescent live labelling of the cellular organelles. Panels (A-E) show the elemental distribution within single whole breast cancer cells reported in Figure 2 of the manuscript showing. **(A)** Cryo-FLM image. Cryo SR-XRF-N images of iridium **(B)**, terbium **(C)**, gadolinium **(D)**. An unambiguous co-localization of gadolinium **(D)**, and terbium **(C)** within a single lysosome labeled by Lysotracker Deep red probe in the cell cytoplasmic extension along with mitochondria depicted by the Ir-Mito probe **(E)**. **Lower panels (F-L)** are images of a MDA-MB-231 cell exposed to $GdTbF_3$-PEG nanohybrids and co-registered to cryo-FLM images. **(F)** Cryo-FLM image. Cryo SR-XRF-N images of zinc **(H)**, iridium **(I)**, gadolinium **(J)**, terbium**(K)** were obtained in that case with a 20 ms dwell-time. **(L)** The corresponding cryo-FLM using computational clearing processing and maximum integrated intensity of image stack (MIP) shows cell nucleus in blue, mitochondria in green and lysosomes in pink **(G)**. Some lamellar structured into mitochondria are again visualized through the iridium elemental map that co-localized with the luminescent signal of the Ir-Mito probe. Registration and merging of the images using the blue channel from the Hoechst cell nucleus labelling and the Zn XRF map allow to observe some co-localization between Gd and Tb XRF map and the lysosomes (cryo-FLM pink channel).

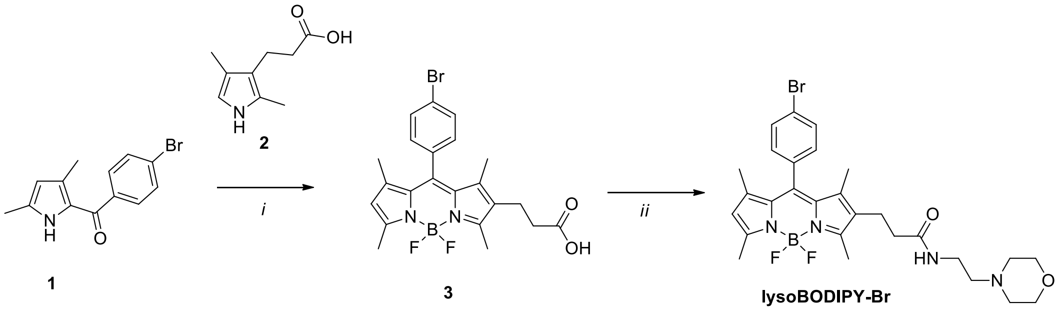


**Figure S5. Synthetic steps for the preparation of lysoBODIPY-Br.** Reagent and conditions: *i)* $POCl_3$, DCM, 3 h; *ii)* TBTU, $NEt_3$, DCM, 16 h. The substrates **1** [ref. 1] and **2** [ref. 2] were prepared according to previously reported procedures. **3** (50 mg, 0.11 mmol) was combined with 4-(2-aminoethyl)morpholine (17 µL, 0.13 mmol) and 2-(1H-benzotriazole-1-yl)-1,1,3,3-tetramethylaminium tetrafluoroborate (TBTU, 1.2 eq.) in dichloromethane (6 mL). Triethylamine (2 eq.) was added and the solution was stirred at room temperature overnight. The solvent was removed under reduced pressure to afford the crude product, which was purified by column chromatography (silica gel, dichloromethane/methanol gradient) to yield the target compound as a bright orange solid (33 mg, 53%).
